# A method for defining tissue injury criteria reveals ligament deformation thresholds are multimodal

**DOI:** 10.1101/2023.01.31.526318

**Authors:** Callan M. Luetkemeyer, Corey P. Neu, Sarah Calve

## Abstract

Soft tissue injuries (such as ligament, tendon, and meniscus tears) are the result of extracellular matrix damage from excessive tissue stretching. Deformation thresholds for soft tissues, however, remain largely unknown due to a lack of methods that can measure and compare the spatially heterogeneous damage and deformation that occurs in these materials. Here, we propose a method for defining *tissue injury criteria*: multimodal strain limits for biological tissues analogous to yield criteria that exist for crystalline materials. Specifically, we developed a method for defining injury criteria for mechanically-driven fibrillar collagen denaturation in soft tissues, using regional multimodal deformation and damage data. We established this new method using the murine medial collateral ligament (MCL) as our model tissue. Our findings revealed that multiple modes of deformation contribute to collagen denaturation in the murine MCL, contrary to the common assumption that collagen damage is driven by strain in the fiber direction alone. Remarkably, our results indicated that hydrostatic strain, or volumetric expansion, may be the best predictor of mechanically-driven collagen denaturation in ligament tissue, suggesting crosslink-mediated stress transfer plays a role in molecular damage accumulation. This work demonstrates that collagen denaturation can be driven by multiple modes of deformation and provides a method for defining deformation thresholds, or injury criteria, from spatially heterogeneous data.

## 1. Introduction

Injuries to soft connective tissues are often debilitative and costly [4, 7]. For example, anterior cruciate ligament tears typically require surgical reconstruction and can lead to early onset osteoarthritis in the knee [32]. Similarly, back pain caused by intervertebral disc injuries is commonly treated by spinal fusion surgery, alleviating pain but eliminating spinal flexibility [13]. Beyond orthopedics, about 1 in 3 women experience a uterosacral ligament tear in their lifetime, which can result in uterine prolapse and require surgical intervention [34].

These injuries are the result of mechanical damage to connective tissue microstructure, which consists of macromolecules (e.g., collagens, elastin, proteoglycans, glycosaminoglycans) collectively referred to as the extracellular matrix (ECM) [23]. Recent evidence suggests that many soft tissue failures result from repeated sub-failure microdamage to the ECM, rather than single overload events [8, 10, 12, 24, 35]. Yet, methods for characterizing the relationship between deformation and tissue microdamage are lacking. In this paper, we propose a combination of experimental and computational methods for defining *tissue injury criteria*, or multimodal mechanical thresholds for soft tissues, using the murine medial collateral ligament (MCL) as our model tissue.

The MCL was chosen for development of this method because of the relative simplicity in comparison to other tissues; it is fairly straight and long, and the bony attachments are ideal grip connections for mechanically testing the tissue without clamping it directly. Importantly, the MCL is predominately composed of type I collagen fibers that are aligned along the axis of loading. Type I collagen fibers are major tensile load-bearing structures composed of collagen molecules bound together by enzymatic crosslinks [6, 23, 31]. For this reason, tendons and ligaments are often modeled as transversely isotropic, which enables the material behavior of the MCL to be investigated independently along and orthogonal to fiber direction.

It is still unknown whether multiple modes of tissue-level deformation significantly contribute to damage and injury, or whether a single strain component is a sufficient predictor. It is commonly assumed that only strain along the axis of collagen fibers contributes to failure and damage in soft tissues [17, 35]. However, emerging evidence has revealed that soft tissues often exhibit unexpected and complex mechanical behavior that defies conventional engineering assumptions; unidirectional stretching does not create a state of uniaxial tension, and deformations are not isochoric, or volume-conserving, as previously believed [19, 21]. In the MCL, the preferential alignment of collagen fibers along the axis of loading will enable the examination of the assumption that fiber strain is sufficient to predict collagen damage. Evidence validating or refuting this idea is important for developing computational studies of injury.

Computational engineering tools such as finite element analysis are an ideal platform with which to study injury risk and prevention. In contrast to experimental studies, computational models can explore all potential variables (e.g., kinematic, anatomic) in a systematic and deterministic way. However, to be quantitatively predictive, these models require accurate, experimentally-determined descriptions of the mechanical behavior of the tissue. To this end, progress has been made to model the multimodal stress-deformation relationship for soft tissues [1, 21, 25]. Still, methods for defining deformation-damage relationships, or injury criteria, are needed to predict injury.

Beyond computational modeling, injury criteria are also essential for quantifying how material differences (e.g., ECM composition and organization) due to age, sex, disease, and degeneration affect injury risk. Understanding how material properties like injury criteria differ between groups will help identify mechanical and therapeutic targets for injury treatment and prevention efforts.

However, the identification of material properties is complicated by unavoidable spatial heterogeneity in soft tissue deformation and damage [2, 19, 21, 24, 35]. To characterize true material properties, we must account for spatial heterogeneity in order to disentangle geometric effects from the material response. Thus, in this work, we employ full-field (i.e., image-based) methods to measure the heterogeneous deformation and damage fields that arise during/after unidirectional stretching of the MCL.

The objectives of this study were to (1) develop a full-field method to identify molecular-level collagen injury criteria for soft tissues and (2) determine whether mechanically-driven collagen denaturation in the MCL is driven by multimodal or unimodal deformation. Findings revealed that both in-plane normal strains (along and orthogonal to collagen fibers) were significant predictors of collagen denaturation in the wild-type murine MCL. Remarkably, positive hydrostatic strain was the best single predictor of collagen denaturation, which we propose is further evidence of the importance of crosslink-mediated stress transfer in mechanically-driven collagen denaturation. The method and data presented here can be used to study how tissue quality and composition affects injury susceptibility, inform computational mechanics models studying injury risk and prevention, and guide the development of tissue material models that incorporate collagen microdamage.

## 2. Methods

### 2.1 Sample preparation

Four 8-12 week old C57BL/6J mice (Jackson Laboratory) were euthanized via CO_2_ asphyxiation and cervical dislocation, according to protocol #2705 approved by the University of Colorado Boulder IACUC. To prepare MCL samples for mechanical testing, hindlimbs were harvested immediately after euthanization and soft tissues other than the MCL and those interior to the knee capsule (e.g., menisci, anterior cruciate ligament) were removed. Care was taken to remove fascia around the MCL without damaging it. Debrided samples were then stained in 1.5 mL 1 × phosphate buffered saline (PBS) with 1:500 Ghost Dye 780 (Tonbo Biosciences) and 3 drops NucBlue (ThermoFisher) for 3 hours at 4 °C with gentle rocking. Ghost Dye (GD) binds to free amine groups to visualize proteins in the MCL, while NucBlue (NB) stains live cell nuclei, which provided additional image texture for deformation tracking.

Following staining, all soft tissues except the MCL were removed from the knee, and the tibial plateau and most of the foot were removed. The femur and tibia were secured in custom 3D printed mechanical testing grips with heat set polystyrene (Fig. 1). All samples were tested the same day as harvest.

**Figure 1.**
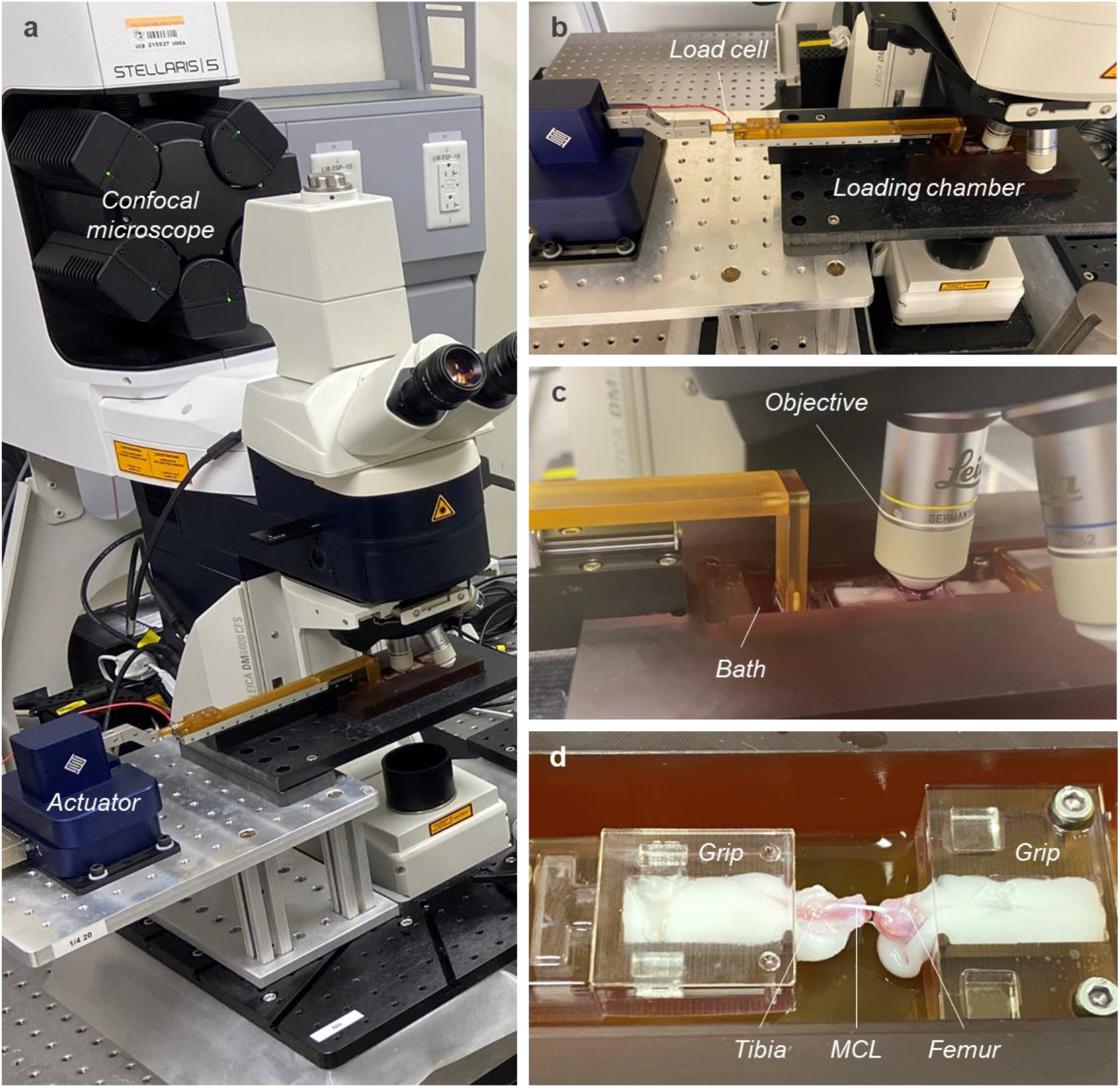
Experimental setup enabled tandem acquisition of image-based deformation and damage measurements. A mechanical testing apparatus was secured to the stage of a confocal microscope (a). The testing apparatus consisted of a mechanical actuator, in-line load cell, and custom loading chamber (b). The loading chamber contained a PBS bath for staining and hydrating the specimen, which was imaged with an immersion objective (c). Custom grips secured the femur and tibia with thermoplastic (d).

### 2.2 Mechanical testing and imaging

With the femur and tibia secured in the custom grips, the grips were attached to a custom mechanical testing rig mounted on an upright confocal microscope (Leica STELLARIS 5) in a bath with 1× PBS (Fig. 1). A FemtoTools micromanipulator was used to drive actuation, and a 1000 g in-line Futek load cell was used to measure forces. A 0.1 N pre-load was used to define the reference state.

With the pre-load set, the PBS bath was drained and replaced with a 5 μM solution of collagen hybridizing peptide (CHP) in 1× PBS. CHP is a molecular probe that binds specifically to denatured collagen molecules [35]. The sample was stained with CHP, *in situ*, for 15 minutes; preliminary testing demonstrated little change in surface CHP staining after 15 minutes. Following baseline staining, the CHP solution was removed and replaced by fresh 1× PBS.

The top surface (~200 μm depth) of the entire MCL was imaged at 1× zoom with a 10× water immersion objective (pixel size of 2.14 μm). Gains and laser intensities for NB and GD channels were set so that the full pixel intensity range (0-255) was observed. For CHP, the gain was adjusted so that the initial pixel intensity range was about 1-100. This ensured that the signal was detectable while allowing for subsequent increases in intensity due to collagen denaturation following mechanical deformation.

With the grip holding the femur fixed to the loading chamber, the grip with the tibia was axially translated by fixed displacements: 500, 750, and 1000 μm. The sample surface was again imaged in each deformed configuration. After each stretch, the sample was returned to the reference state, stained with the CHP solution for 15 minutes, and imaged again. A graphical schematic of the experimental timeline is shown in Figure 2a.

**Figure 2.**
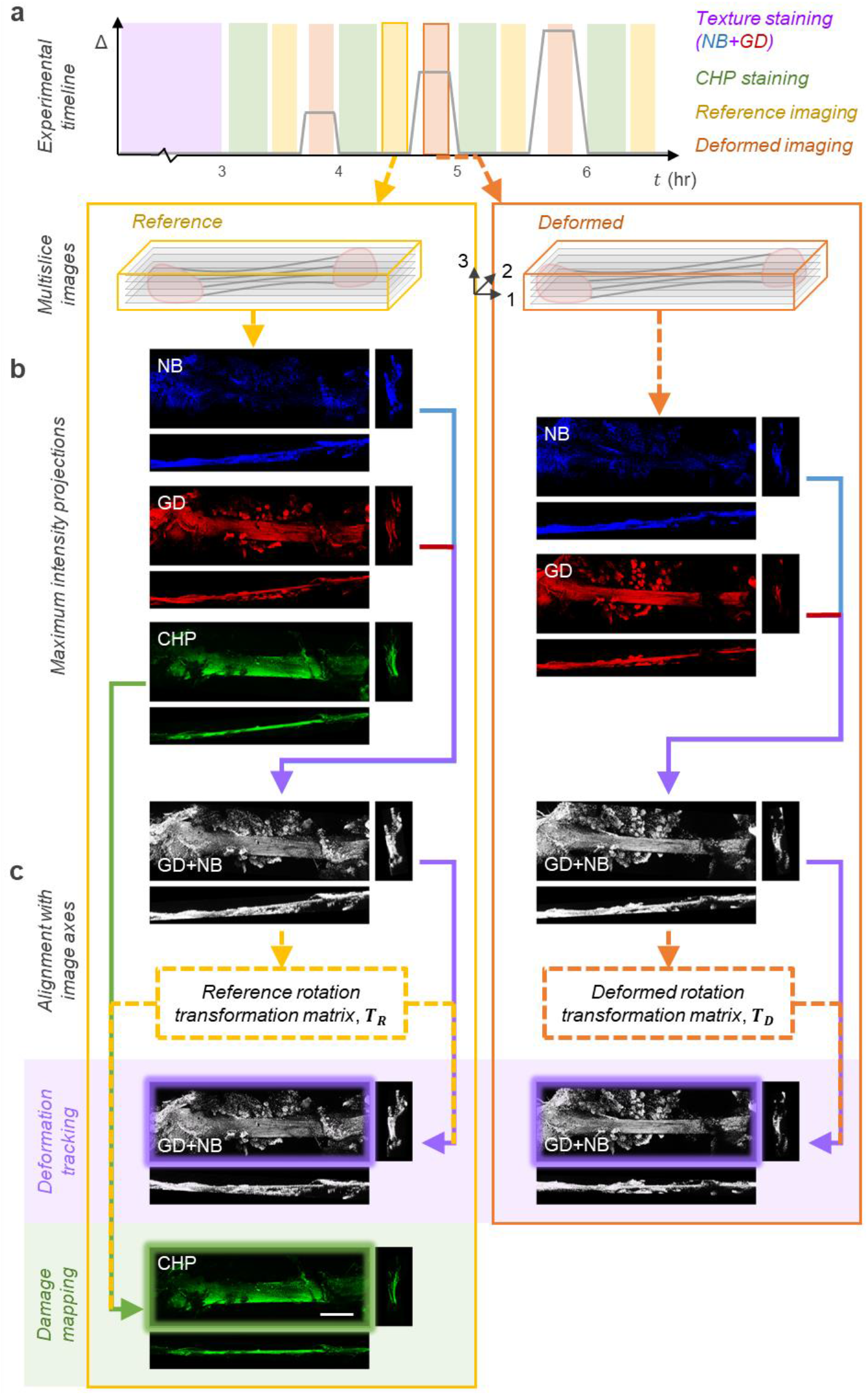
Multislice, multichannel reference/deformed image pairs were pre-processed before image projections were used for injury criteria analysis. The experimental timeline (a) illustrates the order of staining and imaging events in relation to mechanical displacements (Δ) as a function of time (*t*). For each image (reference and deformed), Ghost Dye (GD) and NucBlue (NB) images were combined to create composite images for deformation tracking (b). Rotation transformation matrices were constructed to align the specimen with the global image axes for deformation tracking and damage mapping (c). Scale bar (in final CHP image) represents 1 mm, and all images are shown at the same zoom/magnification.

### 2.3 Image pre-processing

Volume images were opened in ImageJ and saved as .tiff stacks. These multislice images were then read by MATLAB (MathWorks), and blue (NB) and far red (GD) channels were combined, balancing signal from both to ensure both channels were seen in a GD+NB composite image (Fig. 2), which was used for deformation tracking. However, samples rarely demonstrated alignment in the 3, or vertical (*z*), direction throughout the experiments. As a result, surface displacements estimated from *z*-projections of the reference and deformed image stacks were measurably affected. Thus, the built-in function `regionprops3` was used to estimate the principal directions (*θ*_1_, *θ*_2_, *θ*_3_) of the MCL sample in 3D space (relative to the image axes) with a binarized version of each GD+NB image stack. Rotation transformation matrices were then constructed for each reference (***T**_R_*) and deformed (***T**_D_*) image using the product of Euclidean rotation matrices about each axis,

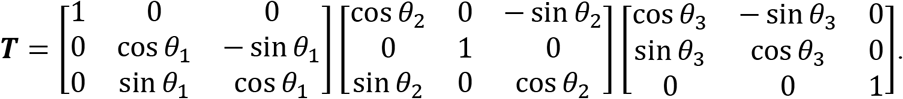

Transformation matrices were used in conjunction with the built-in function `imwarp` to rotate each GD+NB image stack to align with the global coordinate frame. Additionally, for each image in the reference state, the CHP image stack was transformed by the same ***T**_R_*, to provide a 1:1 pixel comparison between deformation and damage fields.

Once rotated and cropped (if necessary, to maintain same size images), maximum intensity *z*-projections of the GD+NB image stacks were obtained for each reference and deformed state and used for deformation tracking. Maximum intensity *z*-projections of the rotated CHP image stacks were prepared for each reference state and used for damage mapping. The alignment and projection steps are summarized in Figure 2. Finally, image masks were constructed for each reference state region of interest (ROI) by manually selecting points along the edge of the MCL, as visualized with the GD+NB *z*-projections.

### 2.4 Deformation mapping

Texture-based displacement tracking proved to be a challenging task for several algorithms tested [3, 9, 14]. Image intensity tended to decrease during stretching, possibly because fewer fluorophores were in any given voxel when large strains were imposed. Thus, the histograms of each deformed image were automatically adjusted to match the respective reference image using `imhistmatch`; this greatly improved automated trackability. Moreover, large rigid body translations and shear concentrations were found to be difficult for many algorithms to track without manual assistance. Thus, displacement fields were first estimated using manually-selected points (e.g., cell nuclei) in each reference-deformed image pair, with the assistance of the built-in MATLAB function `cpselect`. 2D linear interpolation was used to approximate displacement vectors for each pixel location in the ROI from the scattered displacement data at manually-selected points. A 2D moving average filter with a small, anisotropic kernel was used to filter this displacement field (***u***^(*m*)^) before using it to warp the deformed image to nearly match the reference image. The warped deformed image and its reference image were then given to the function `imregdemons` to compute auto-tracked displacement fields for fine-tuning (***u***^(*a*)^). Since both displacement fields were determined in the reference configuration, they were superimposed to produce the final displacement fields,

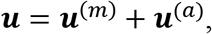

which were used to compute Green-Lagrange strains for injury criteria analysis.

### 2.5 Damage analysis

Binary maps of collagen damage were constructed by applying a threshold to CHP *z*-projections. Because samples were stained multiple times (after each deformation) during experiments, increasing CHP intensity resulting from more stain time (rather than damage) was a concern. Several preliminary tests were run to discern if increased CHP signal due to stain time could be differentiated from that caused by an actual increase in collagen denaturation. Whether the damage was due to mechanical deformation or the application of heat, a similar trend was observed: both the mean and standard deviation of the image intensity histogram changed when damage was induced, but only the mean changed substantially with increasing stain time (Fig. 4a). Thus, to account for intensity changes due to stain time, the threshold was updated for each image,

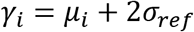

where *γ_i_* is the damage intensity threshold for the *i^th^* image, *μ_i_* is the mean image intensity of the *i^th^* ROI, and *σ_ref_* is the standard deviation of the image intensity histogram for the ROI in the original (baseline) reference image. This equation for the damage intensity threshold accounts for increasing stain time, while resolving changes in CHP intensity due to mechanical stretching (Fig. 4b-c).

**Figure 3.**
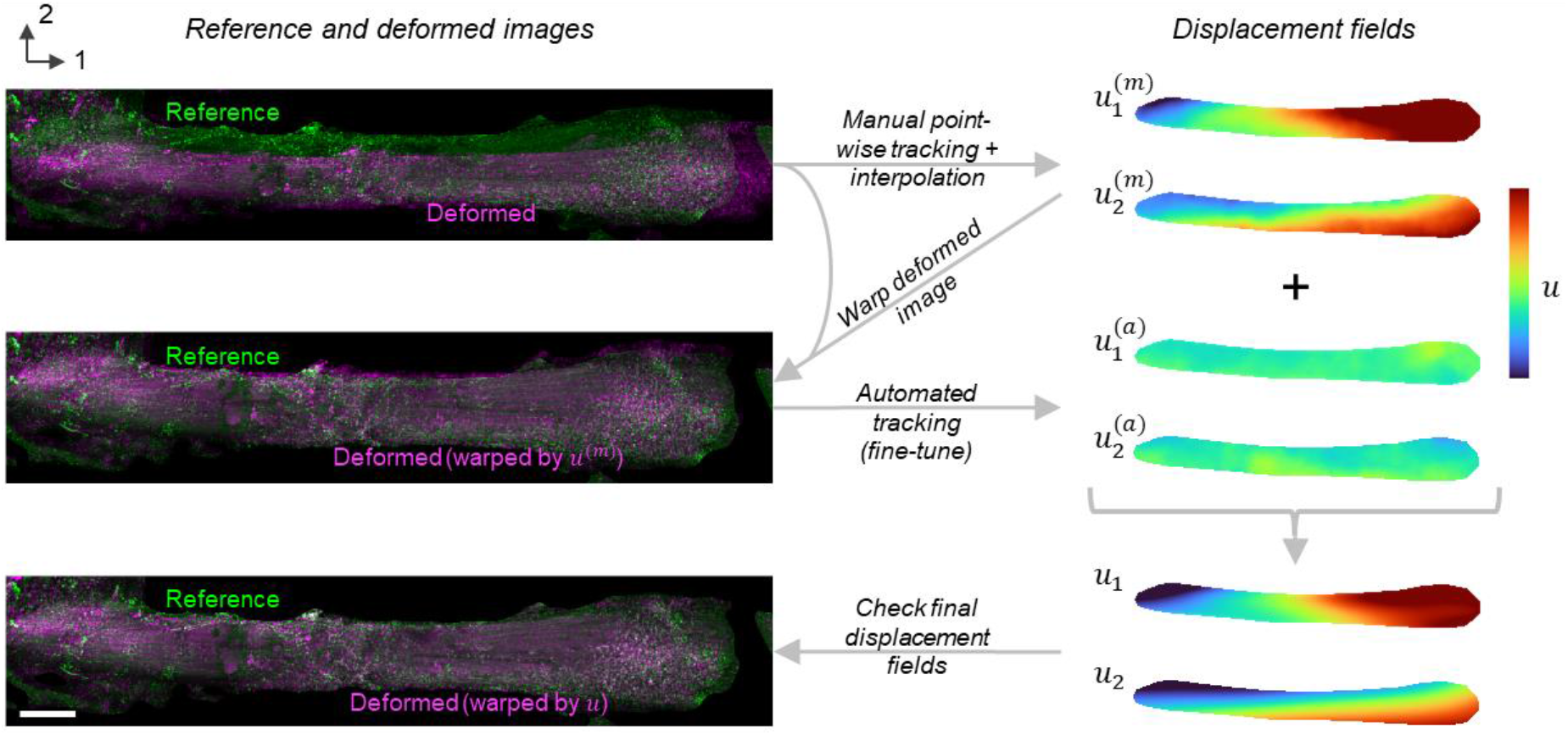
The deformation mapping process consisted of both manual and automated image tracking elements. Briefly, the process entailed (1) manual point-wise tracking and interpolation, (2) warping the image of the deformed configuration by the manually-constructed displacement field, (3) automated deformation tracking between the reference and warped deformed configurations, and (4) superposition of the manually-constructed displacement field (an approximation) with the automatically-tracked displacement field (for fine-tuning). Scale bar represents 500 μm, and all images are shown at the same zoom/magnification.

**Figure 4.**
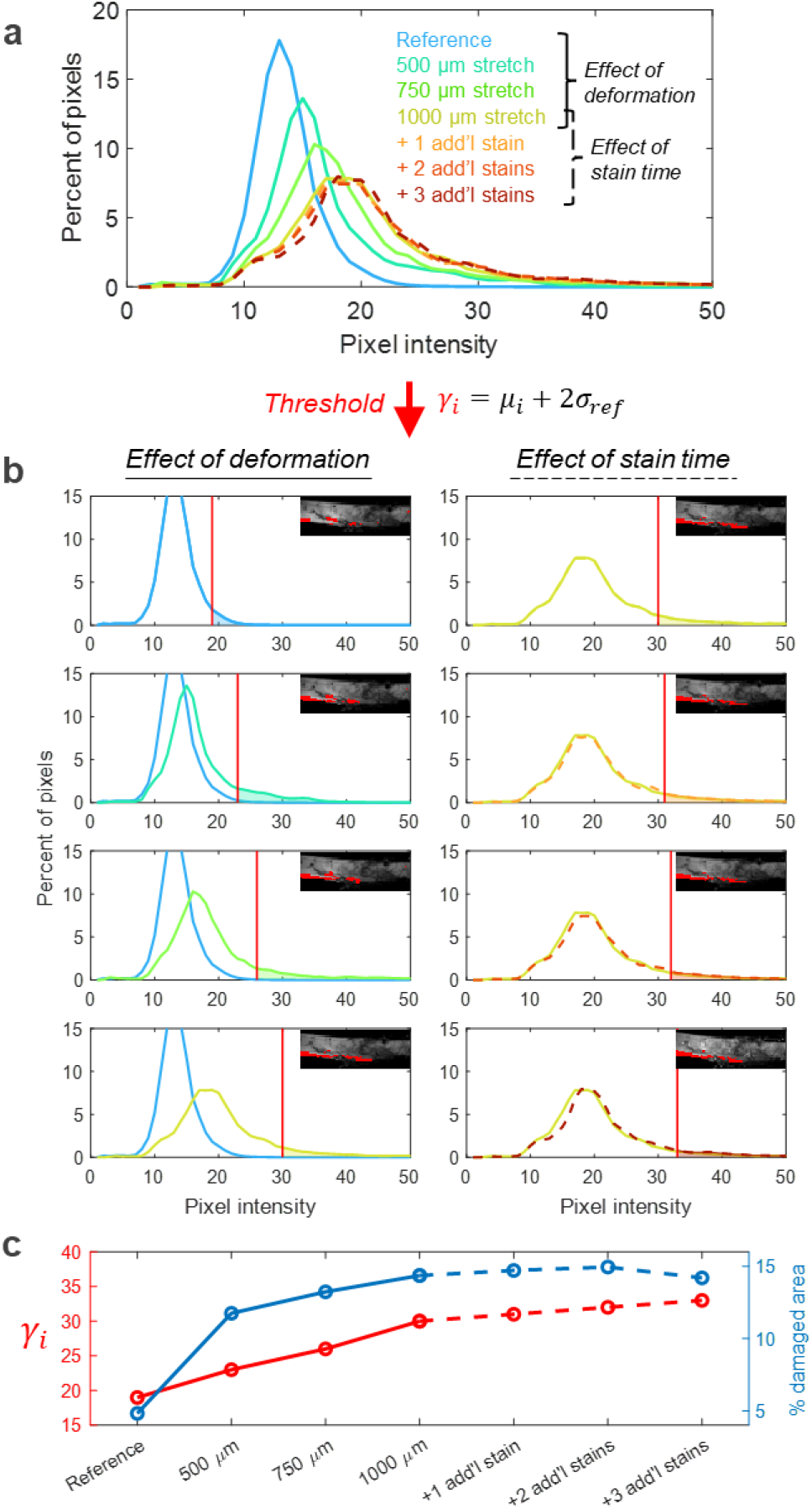
The image intensity damage threshold was tested on a specimen subjected to repeated staining with and without additional mechanical deformation. The CHP image intensity histogram (a) changes with deformation (solid lines, blue to yellow) and stain time (dashed lines, orange to red) in a single specimen. The moving threshold (*γ_i_*), represented by the bright red vertical line (b), accounts for the rightward shifts in mean intensity due to stain time (right). The region identified with significant collagen denaturation (red pixels in images over histograms) grew with increasing deformation, but stagnated with increasing stain time alone (blue line, c).

### 2.6 Construction of collagen injury criteria for individual samples

2D Green-Lagrange strain fields were calculated from the estimated displacement fields,

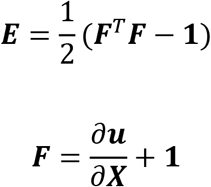

Where ***F*** is the deformation gradient tensor, **1** is the identity tensor, and ***X*** is the coordinate vector in the reference configuration. From the strain tensor (***E***), the in-plane principal strain fields were computed,

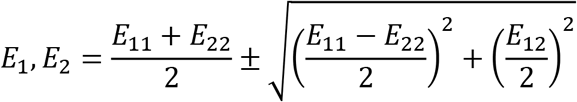

as well as the hydrostatic (*E_hyd_*), deviatoric (***E’***), and equivalent (*E_eq_*) strains:

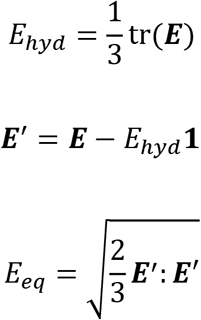

Strain and damage fields from each deformation state were downsized to 10% of the original size to reduce noise and yield more manageable datasets. Each pixel in the downsampled strain and damage fields was considered a single data point, with the in-plane normal (*E_11_* and *E_22_*) and shear (E_12_) strains regarded as independent predictor variables, and collagen damage as a binary dependent variable. Because deformation is heterogeneous, some material points experienced deformation that far exceeded the injury criterion (damage threshold), while others remained far below the threshold. The objective was to identify the best-fit boundary between these two populations of points – damaged and not damaged – which are scattered in strain space. Here, this was achieved with logistic regression. Because most datasets contain fewer “damage” data points than “no damage”, the classification analysis is skewed by imbalanced classifiers if all data points are used. Thus, a random sampling of data points from the larger group (equal to the number of points in the smaller group) was used to balance the number of classifiers from both populations. This process is illustrated for a single sample in Figure 5.

**Figure 5.**
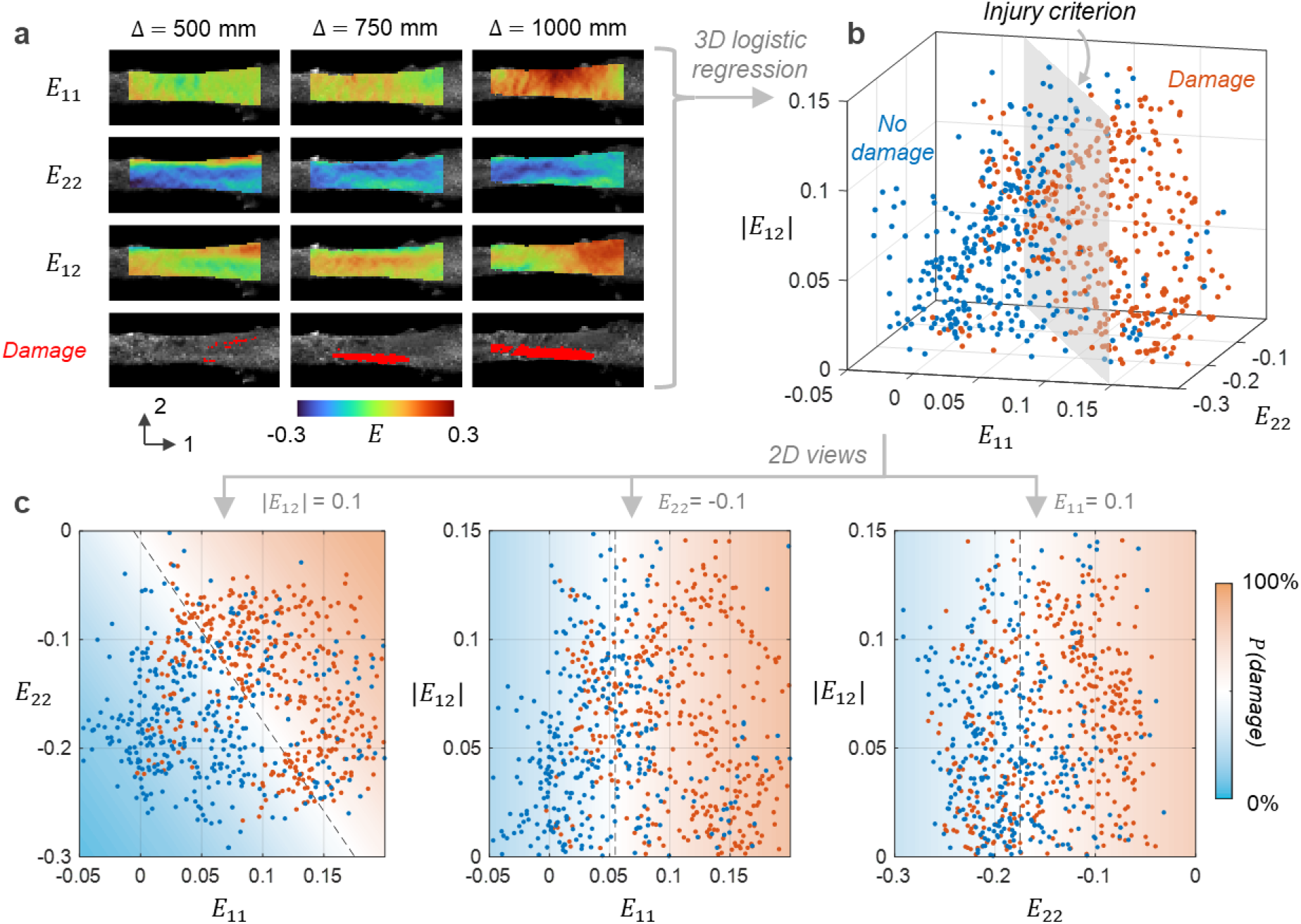
An initial injury criterion was constructed with spatially heterogeneous data from a single experiment. In-plane strain and damage fields from three deformation (Δ) states (a) generated many deformation-damage data points in a 3D strain space (b). Logistic regression was used to determine an injury criterion, or best-fit strain-based boundary between damaged and undamaged points. 2D slices facilitate visualization of the strain space and identification of the strain components that are significant predictors of collagen denaturation (c). The regression-derived probability of damage is shown with a red-blue colormap.

Because the in-plane Green-Lagrange strains depend on the global coordinate frame, injury criteria were also constructed using two sets of frame-indifferent measures of strain: hydrostatic and equivalent (i.e., von Mises) strains and in-plane principal strains. These analyses were conducted using the same data points.

### 2.7 Statistical analyses

Data collected with samples from three different animals were combined to determine injury criteria for the wild-type murine MCL. For each analysis – in-plane strains, hydrostatic-equivalent strains, and principal strains – logistic regression was performed with the function mrnfit. Strain components with regression coefficient p-values less than 0.05 were considered to be significant predictors of collagen denaturation.

## 3. Results

Injury criteria constructed with deformation-damage data combined from three specimens indicate that collagen denaturation can be driven by multiple modes of tissue-level deformation (Fig. 6). 3D logistic regression analysis of in-plane strains (Fig. 6a) revealed that both in-plane normal strains (*E*_11_ strain in the fiber direction, and *E*_22_, strain orthogonal to the fibers) were significant predictors of collagen denaturation (p < 0.001), while shear strain (*E*_12_) was not (p > 0.05).

**Figure 6.**
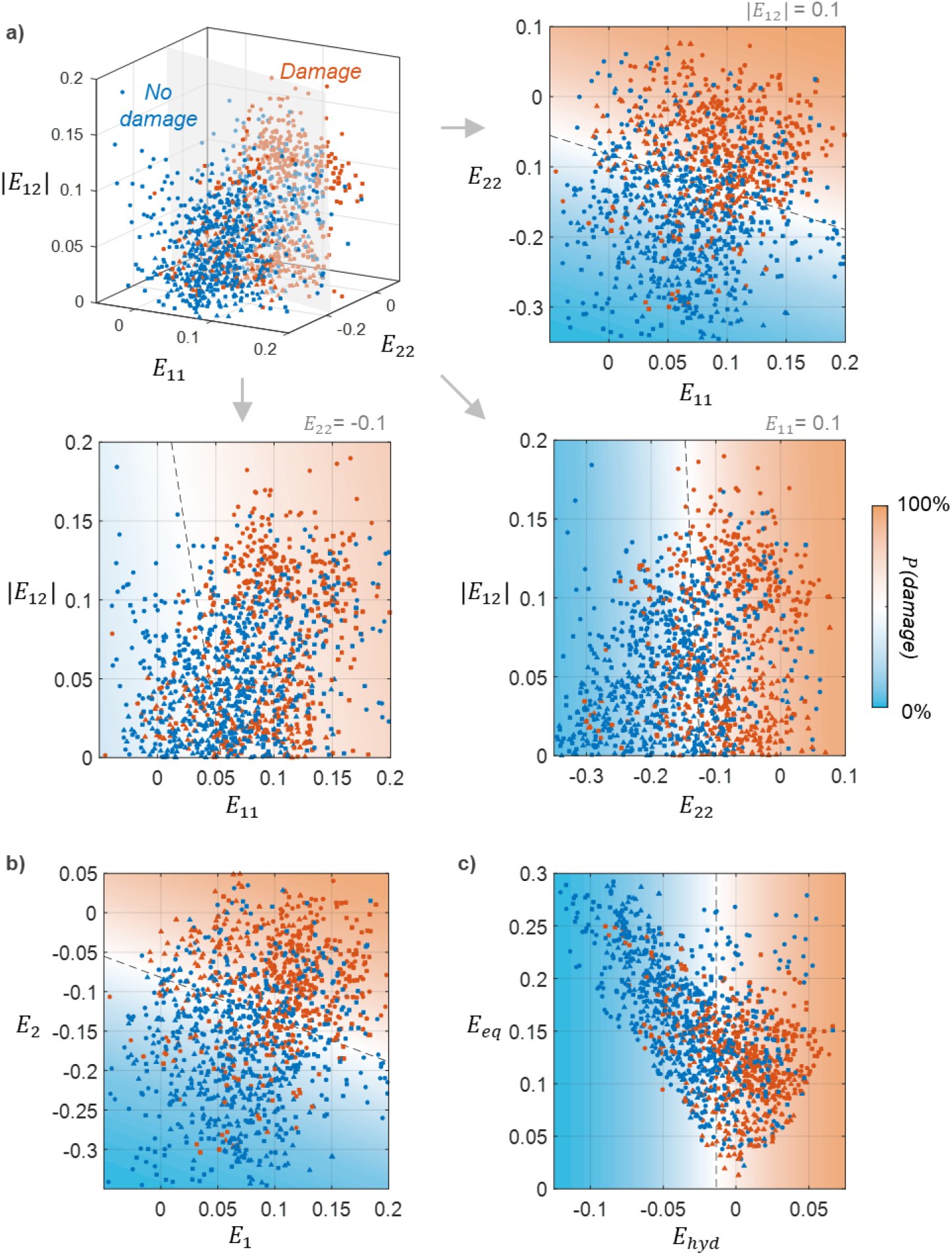
Injury criteria for murine MCLs indicate collagen denaturation is driven by multimodal tissue-level deformation. The same data points were used to construct injury criteria in terms of in-plane surface strains (a), in-plane principal strains (b), and equivalent (*E_eq_*)and hydrostatic (*E_hyd_*) strains (c). Regression analyses indicate that both in-plane normal strains, hydrostatic strain, and both in-plane principal strains were significant predictors of collagen denaturation (p < 0.001), while shear and equivalent strains were not (p > 0.05). Data from biological replicates (n = 3) are plotted with different symbols (circle, triangle, or square).

Injury criteria constructed in terms of frame-indifferent measures of strain supported the result that multiple modes of strain drive collagen denaturation. Both in-plane principal strains were significant predictors of collagen damage (p < 0.001), confirming that both normal strains contribute to collagen damage. This result prompted the exploration of hydrostatic strain as a driver of collagen denaturation. 2D logistic regression analysis of the data in terms of hydrostatic and equivalent strains revealed that hydrostatic strain was the single best predictor of collagen denaturation of all the strain measures investigated (p << 0.001). Equivalent strain, however, was not a good predictor (p > 0.05). The vertical threshold in Fig. 6c demonstrates that collagen denaturation depends entirely on hydrostatic strain within the hydrostatic-equivalent strain space. This threshold was identified to be near 0% hydrostatic strain, with a 95% confidence interval of (−9.5%, 6.8%).

## 4. Discussion

This study aimed to develop a method to identify injury criteria, or multimodal deformation thresholds, for fibrillar collagen in soft tissues using the murine MCL as a model example. The full-field method created takes advantage of spatially heterogeneous deformation and damage by collecting many data points throughout a multimodal strain space with a single experiment. Using data from just three specimens, logistic regression analyses were able to determine that multiple modes of strain were statistically significant predictors of collagen denaturation. Surprisingly, hydrostatic strain (a multimodal deformation measure) was the single best predictor of damage. This finding has important implications on our understanding of the mechanisms governing deformation-induced collagen denaturation as well as computational modeling of soft tissue injury. Additionally, the method developed will enable future studies to examine the relationship between tissue composition and injury risk.

### 4.1 Further evidence for crosslink-mediated collagen denaturation

Prior to this study, it has been widely assumed that only strain in the direction of collagen fibers would contribute to collagen damage [17, 35]. Our findings confirm that tissue-level strain in the direction of collagen fibers is a significant predictor of collagen denaturation (p < 0.001). However, our results also indicate that strain orthogonal to the fibers significantly contributes to molecular damage (p < 0.001). Analysis in terms of in-plane principal strains confirmed these results.

The finding that both in-plane normal strains are significant predictors of collagen damage led to the question: can volumetric expansion drive collagen denaturation? Unexpectedly, we found that hydrostatic strain was the single best predictor of collagen denaturation, while equivalent (i.e., von Mises) strain, which is often used to predict failure in ductile materials, was not a significant predictor (p > 0.05). This result is the opposite of what classic anisotropic yield criteria like Hill’s generalized criterion or the Logan-Hosford criterion would predict [11, 18]. Moreover, we consistently found the hydrostatic strain threshold to be about 0%, suggesting that volumetric expansion, rather than fiber extension alone, contributes to mechanically-driven collagen denaturation. In other words, the extension of collagen fibers alone does not necessarily lead to collagen molecule denaturation if the tissue laterally contracts enough to avoid a local increase in volume.

Our surprising result that hydrostatic deformation contributes to denaturation provides further evidence for the hypothesis that mechanical failure of collagen molecules is driven by crosslink-mediated stress transfer, as proposed by Zitnay and colleagues [35]. As collagen fibers stretch, collagen triple helices are known to slide relative to each other, with stress transferred between molecules via crosslinks bound to single α-chains [6]. The force required to rupture individual covalent crosslinks was previously estimated to be an order of magnitude greater (6-8 nN) [6] than that required to pull a single α-chain out of a collagen molecule [35]. Taken together with our current finding that collagen denaturation occurs in regions with positive hydrostatic strain, we propose that crosslink-mediated lateral tensile forces play a large role in mechanically-driven collagen denaturation (Fig. 7). Positive lateral stresses often occur in tissues, even when loaded uniaxially; our previous work demonstrated that fiber splay (the tendency of fibers to spread out near attachments) can cause lateral expansion in a uniaxial loaded ligament [20, 21, 22].

**Figure 7.**
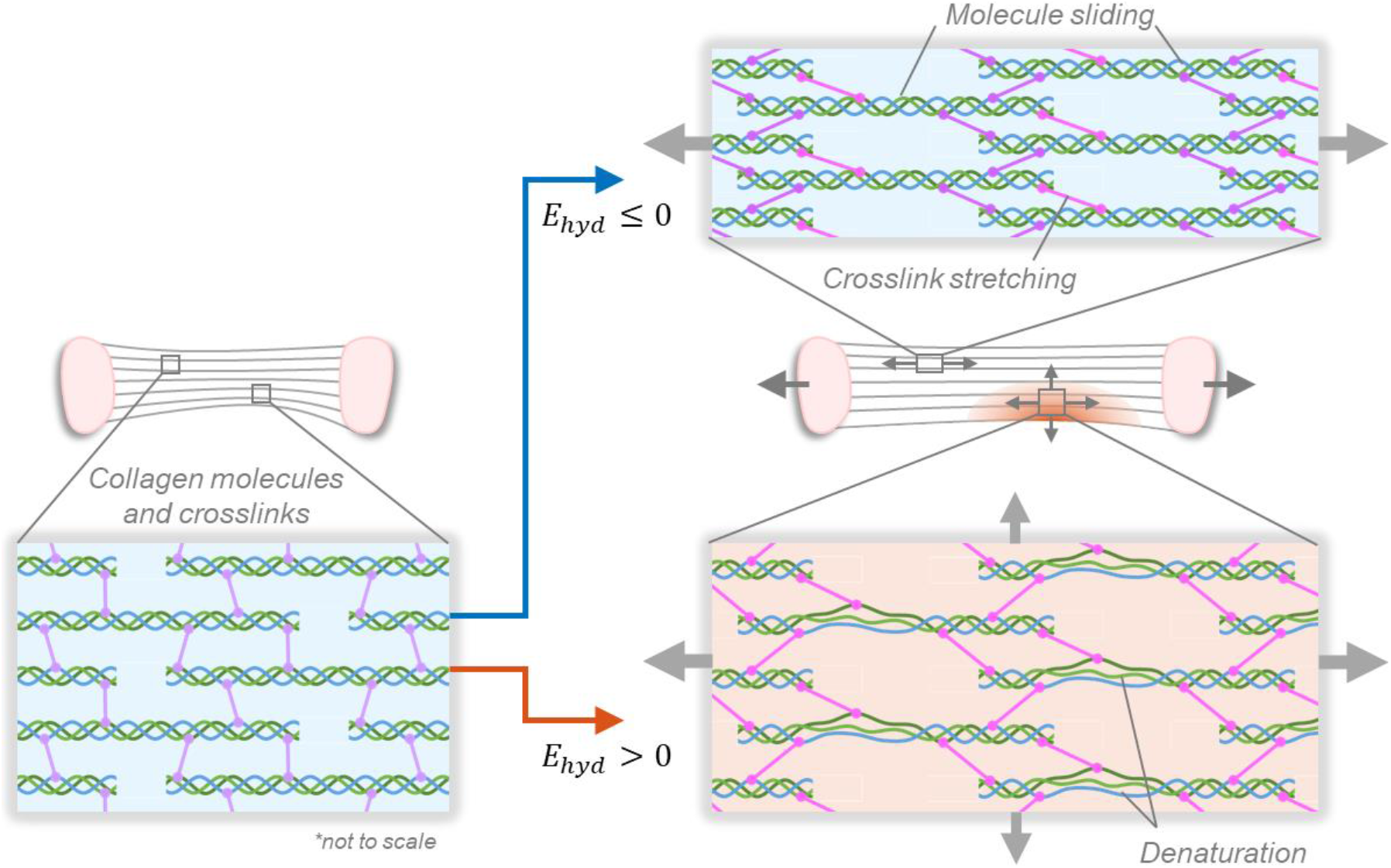
We hypothesize that collagen crosslinks drive mechanically-driven collagen denaturation through lateral force transfer. Our findings reveal that ligaments can withstand more strain along the primary fiber direction (*E*_11_) without sustaining significant collagen denaturation if allowed to laterally contract. Regions that experienced volumetric expansion (*E_hyd_* > 0) due to less lateral contraction (*E*_22_) were significantly more likely to incur collagen damage than regions that did not. Thus, we propose that lateral forces transmitted via molecular crosslinks are primarily responsible for the mechanical unwinding of collagen triple helices.

While a thorough investigation into the molecular mechanisms at play is outside the scope of this work, we anticipate that this study will motivate deeper inspection of the role of molecular crosslinks in soft tissue injury.

### 4.2 Broader implications

The results of this study are critical for informing computational and theoretical models of injury. Our findings indicate that multiple modes of tissue-level deformation contribute to collagen molecule damage, even when collagen fibers are highly aligned in one direction. Thus, studies employing finite element models to investigate questions about soft tissue injury need to consider more than fiber direction stretch when deciding whether predicted deformation patterns will lead to damage. The data and analysis presented here will also guide the development of more comprehensive material models that incorporate sub-failure damage as an internal variable [26, 28], further advancing computational research on soft tissue injuries.

Moreover, the method developed will be used for hypothesis-driven studies on the role of tissue composition and organization on injury risk. Just as yield criteria for crystalline materials are characterized and compared, this method allows a materials-based comparison of heterogeneously deforming soft tissues. For example, injury criteria can used to investigate the effect of age or sex-related differences on injury susceptibility, where group disparities in macroscopic mechanical behavior can result from geometric (rather than constitutive) differences. This full-field method has the potential to quantitatively distinguish between the effects of material and anatomical disparities.

### 4.3 Limitations and future work

While the findings presented here show a significant effect of multiple modes of strain on collagen denaturation, they should be viewed within the context of the study’s limitations. Most notably, this work is limited to 2D information and analysis, since confocal microscopy cannot resolve images deep within a live, ECM-rich tissue. The lack of 3D information means we had to project a 6D strain space into a 3D (*E*_11_, *E*_22_, and *E*_12_) one. This likely contributed to the scatter in injury criteria data (i.e., the misclassified “damage” or “no damage” points). Moreover, plane strain (*E*_33_ = 0) was assumed to compute equivalent and hydrostatic strains; however, without out-of-plane deformation and damage data, we cannot be certain of the true hydrostatic strain threshold. Nevertheless, even without this information, results were consistent both within and across biological replicates.

We also note that while our full-field data provided many observations throughout the surface strain space, the entire strain space was not systematically probed. Data is limited to the strain states that arise during unidirectional loading. Hence, even though we did not find tissue-level shear strain to be a significant predictor of collagen denaturation in this study, it is possible that a dependence might be found if larger shear strains were experimentally imposed.

Moreover, the method presented here greatly simplified damage analysis using logistic regression. Damage was considered a binary variable; each material point was labeled as either having damage (1), or not (0). In reality, however, each pixel in a CHP channel image contains numerous collagen molecules, some of which are denatured and bound to CHP, while others are undamaged. A more accurate way to model collagen damage from CHP images will be to allow damage to be a continuous variable, taking on fractional values that represent the proportion of molecules with damage. Further, logistic regression limited our analysis to linear thresholds. Future work will aim to resolve both of these shortcomings.

Additionally, we considered only the strain state as a causal variable, as it was directly measurable from images. The stress state is related, but computing the heterogeneous stress field requires assumptions about the constitutive (i.e., stress-strain) relationship. It is unknown whether modifications to the tissue microstructure or composition (e.g., with damage, disease, age) will alter strain or stress-based thresholds. We plan to examine this question in future studies.

In this work, we focused on collagen denaturation, which was spatially mapped using CHP. However, soft tissues are comprised of many ECM components [30], and it is probable that cells and other ECM components have independent injury criteria. In future work, we intend to use other image-based techniques [8, 15, 24] to map ECM and cellular damage to construct a more comprehensive view of soft tissue injury criteria.

Finally, the length scale of the data used in this study is an order of magnitude smaller than what non-invasive, *in vivo* imaging modalities like MRI and ultrasound can achieve: tens vs. hundreds of microns, respectively. However, the pixels used for analysis were larger than that of a single murine collagen fiber [29, 33], so it is possible that the criteria developed here would hold at larger length scales. If so, injury criteria have the potential to inform the use of full-field tissue deformation measured with MRI [16, 19, 21] or ultrasound [5, 27] as a biomarker for early stages of tissue injury and degeneration. We plan to explore this idea in future work.

## 5. Conclusion

In this study, we developed the first full-field method for defining soft tissue injury criteria: multimodal strain thresholds analogous to yield criteria. Using the murine MCL as a model tissue, our findings revealed that multiple modes of deformation are significant predictors of collagen denaturation, which challenges the conventional assumption that only strain along the axis of collagen fibers contributes to damage. Positive hydrostatic strain was the single best predictor of collagen denaturation in this study, which contrasts with the predictions of classic yield criteria and suggests that crosslink-mediated stress transfer plays an important role in mechanically-driven collagen denaturation. The method we developed will be a valuable tool for understanding the role of these collagen crosslinks and other ECM constituents in injury susceptibility, as well as for informing computational models of soft tissue injury.

## Acknowledgements

The authors would like to thank Catalina Bastías, Dr. Stephanie Ellyse Schneider, Emily Miller, Dr. Virginia Ferguson, Dr. Nancy Emery, Dr. Michael Thouless, and Dr. Ellen Arruda for helpful discussions and suggestions. CML was supported by Schmidt Science Fellows, in partnership with the Rhodes Trust. This work was additionally supported by NIH DP2 AT009833 (SC) and R01 AR063712 (CPN).

## References

1. Avril, Stéphane, and Sam Evans, eds. Material parameter identification and inverse problems in soft tissue biomechanics. Cham, Switzerland: Springer International Publishing, 2017.

2. Barreto, Isabella Silva, et al. “Nanoscale characterization of collagen structural responses to in situ loading in rat Achilles tendons.” Matrix Biology 115 (2023): 32–47.

3. Blaber, J., B. Adair, and Antonia Antoniou. “Ncorr: open-source 2D digital image correlation matlab software.” Experimental Mechanics 55.6 (2015): 1105–1122.

4. Choi, Kyung-Chul, et al. “Cost-effectiveness of microdiscectomy versus endoscopic discectomy for lumbar disc herniation.” The Spine Journal 19.7 (2019): 1162–1169.

5. Couppé, Christian, et al. “Ultrasound speckle tracking of Achilles tendon in individuals with unilateral tendinopathy: a pilot study.” European Journal of Applied Physiology 120 (2020): 579–589.

6. Depalle, Baptiste, et al. “Influence of cross-link structure, density and mechanical properties in the mesoscale deformation mechanisms of collagen fibrils.” Journal of the Mechanical Behavior of Biomedical Materials 52 (2015): 1–13.

7. Filbay, Stephanie Rose, et al. “Long-Term quality of life, work limitation, physical activity, economic cost and disease burden following ACL and meniscal injury: a systematic review and meta-analysis for the OPTIKNEE consensus.” British Journal of Sports Medicine 56.24 (2022): 1465–1474.

8. Fung, David T., et al. “Second harmonic generation imaging and Fourier transform spectral analysis reveal damage in fatigue-loaded tendons.” Annals of Biomedical Engineering 38.5 (2010): 1741–1751.

9. Ghosh, Soham, et al. “Deformation microscopy for dynamic intracellular and intranuclear mapping of mechanics with high spatiotemporal resolution.” Cell Reports 27.5 (2019): 1607–1620.

10. Gooyers, Chad E., et al. “Characterizing the combined effects of force, repetition and posture on injury pathways and micro-structural damage in isolated functional spinal units from sub-acute-failure magnitudes of cyclic compressive loading.” Clinical Biomechanics 30.9 (2015): 953–959.

11. Hill, Rodney. “Theoretical plasticity of textured aggregates.” Mathematical Proceedings of the Cambridge Philosophical Society. Vol. 85. No. 1. Cambridge University Press, 1979.

12. Iatridis, James C., Jeffrey J. MacLean, and David A. Ryan. “Mechanical damage to the intervertebral disc annulus fibrosus subjected to tensile loading.” Journal of Biomechanics 38.3 (2005): 557–565.

13. Kapetanakis, Stylianos, Nikolaos Gkantsinikoudis, and Georgios Charitoudis. “The role of full-endoscopic lumbar discectomy in surgical treatment of recurrent lumbar disc herniation: a health-related quality of life approach.” Neurospine 16.1 (2019): 96.

14. Landauer, A. K., et al. “A q-factor-based digital image correlation algorithm (qDIC) for resolving finite deformations with degenerate speckle patterns.” Experimental Mechanics 58 (2018): 815–830.

15. Lane, Brooks A., et al. “Targeted gold nanoparticles as an indicator of mechanical damage in an elastase model of aortic aneurysm.” Annals of Biomedical Engineering 48.8 (2020): 2268–2278.

16. Lee, Woowon, et al. “High frame rate deformation analysis of knee cartilage by spiral dualMRI and relaxation mapping.” Magnetic Resonance in Medicine 89.2 (2023): 694–709.

17. Lin, Allen H., et al. “Collagen fibrils from both positional and energy-storing tendons exhibit increased amounts of denatured collagen when stretched beyond the yield point.” Acta Biomaterialia (2022) In press.

18. Logan, Roger W., and William F. Hosford. “Upper-bound anisotropic yield locus calculations assuming< 111>-pencil glide.” International Journal of Mechanical Sciences 22.7 (1980): 419–430.

19. Luetkemeyer, Callan M., et al. “Full-volume displacement mapping of anterior cruciate ligament bundles with dualMRI.” Extreme Mechanics Letters 19 (2018): 7–14.

20. Luetkemeyer, Callan M., et al. “Fiber splay precludes the direct identification of ligament material properties: Implications for ACL graft selection.” Journal of Biomechanics 113 (2020): 110104.

21. Luetkemeyer, Callan M., et al. “Constitutive modeling of the anterior cruciate ligament bundles and patellar tendon with full-field methods.” Journal of the Mechanics and Physics of Solids 156 (2021): 104577.

22. Mallett, Kaitlyn F., and Ellen M. Arruda. “Digital image correlation-aided mechanical characterization of the anteromedial and posterolateral bundles of the anterior cruciate ligament.” Acta Biomaterialia 56 (2017): 44–57.

23. McKee, Turney J., et al. “Extracellular matrix composition of connective tissues: a systematic review and meta-analysis.” Scientific Reports 9.1 (2019): 1–15.

24. Provenzano, Paolo P., et al. “Subfailure damage in ligament: a structural and cellular evaluation.” Journal of Applied Physiology 92.1 (2002): 362–371.

25. Raghupathy, R., et al. “Identification of Regional Mechanical Anisotropy in Soft Tissue Analogs.” Journal of Biomechanical Engineering. 133.9 (2011): 091011.

26. Rausch, M. K., G. E. Karniadakis, and J. D. Humphrey. “Modeling soft tissue damage and failure using a combined particle/continuum approach.” Biomechanics and Modeling in Mechanobiology 16.1 (2017): 249–261.

27. Rebholz, Brandon, Fei Zheng, and Mohamed Almekkawy. “Two-dimensional iterative projection method for subsample speckle tracking of ultrasound images.” Medical & Biological Engineering & Computing 58 (2020): 2937–2951.

28. Simo, Juan C., and J.W. Ju. “Strain-and stress-based continuum damage models—I. Formulation.” International Journal of Solids and Structures 23.7 (1987): 821–840.

29. Stevenson, Karen, et al. “Functional changes in bladder tissue from type III collagen-deficient mice.” Molecular and Cellular Biochemistry 283 (2006): 107–114.

30. Tonti, Olivia R., et al. “Tissue-specific parameters for the design of ECM-mimetic biomaterials.” Acta Biomaterialia 132 (2021): 83–102.

31. Vallet, Sylvain D., and Sylvie Ricard-Blum. “Lysyl oxidases: from enzyme activity to extracellular matrix cross-links.” Essays in Biochemistry 63.3 (2019): 349–364.

32. Webster, Kate E., and Timothy E. Hewett. “Anterior cruciate ligament injury and knee osteoarthritis: an umbrella systematic review and meta-analysis.” Clinical Journal of Sport Medicine 32.2 (2022): 145–152.

33. Weis, Sara M., et al. “Myocardial mechanics and collagen structure in the osteogenesis imperfecta murine (oim).” Circulation Research 87.8 (2000): 663–669.

34. Zhu, Ya-pei, et al. “Evaluation of extracellular matrix protein expression and apoptosis in the uterosacral ligaments of patients with or without pelvic organ prolapse.” International Urogynecology Journal 32.8 (2021): 2273–2281.

35. Zitnay, Jared L., et al. “Molecular level detection and localization of mechanical damage in collagen enabled by collagen hybridizing peptides.” Nature Communications 8.1 (2017): 1–12.

